# Deep conservation of mitochondrial HSP60 structure with lineage-specific and context-dependent regulation reflects thermal resilience in cnidarians

**DOI:** 10.1101/2025.10.29.685423

**Authors:** Saborni Chowdhury, Avery J. Kruger, Liza M. Roger

## Abstract

Heat shock proteins (HSPs) are ubiquitous molecular chaperones that safeguard proteostasis under stress. We first investigated the expression dynamics of the mitochondrial chaperonin HSP60 across diverse cnidarians to understand its stress-responsive regulation. Using immunoblotting, we quantified HSP60 expression in *Pocillopora acuta* (reef-building coral), *Exaiptasia diaphana* (sea anemone), and *Cassiopea xamachana* (upside-down jellyfish). In *P. acuta*, HSP60 was not detected at the fragment scale under either control or heat stress, whereas isolated cells exhibited transient HSP60 expression during exposure to both control and heated temperatures (+5 °C above optimum), indicating that HSP60 regulation in this coral is strongly context-dependent and potentially suppressed at the tissue level. In contrast, *E. diaphana* and *C. xamachana* showed gradual, and temperature-dependent accumulation of HSP60 over 24 h under heated conditions (+5 °C above its thermal optimum), however *C. xamachana* also displayed constitutive basal expression under control conditions. These contrasting profiles highlight clear lineage-specific differences in HSP60 regulation among cnidarians. The consistent antibody cross-reactivity observed across all three species then prompted us to explore the evolutionary basis of this conservation. Phylogenetic analyses of HSP60 sequences confirmed that cnidarian proteins are orthologous to the canonical vertebrate HSP60 (human HSPD1), demonstrating deep structural and evolutionary conservation of this chaperonin across Metazoa. Collectively, these findings reveal that while HSP60 is evolutionarily ancient and conserved, its regulation under thermal stress varies across lineages and physiological context, reflecting complex modulation of mitochondrial proteostasis in early-diverging metazoans. This lineage- and context-dependent regulatory framework provides new insight into how chaperone plasticity contributes to cnidarian thermal tolerance and the differential susceptibility of reef taxa to bleaching under ocean warming.

**Significance Statement:** Heat shock protein 60 (HSP60) is a highly conserved mitochondrial chaperonin critical for maintaining protein homeostasis, yet its regulatory dynamics across early-diverging animal lineages are poorly understood. By first comparing the expression responses of three phylogenetically and ecologically distinct cnidarians—the coral *Pocillopora acuta*, the sea anemone *Exaiptasia diaphana*, and the upside-down jellyfish *Cassiopea xamachana*—we uncovered clear lineage-specific differences in HSP60 regulation. *P. acuta* showed no detectable HSP60 induction in intact tissue, underscoring strong context-dependence that may prevent the deployment of this critical molecular defense mechanism, reflecting its high thermal susceptibility. In contrast, *E. diaphana* and *C. xamachana* displayed gradual, temperature-dependent accumulation aligning with their thermal flexibility, with *C. xamachana* also displaying constitutive basal levels under control condition. The consistent antibody cross-reactivity across all three species then led us to investigate evolutionary conservation, revealing that cnidarian HSP60s are orthologous to the canonical HSP60 (human HSPD1). This demonstrates that HSP60 is deeply conserved from cnidarians to mammals, yet its stress-responsive regulation has diversified across lineages and physiological contexts. This lineage- and context-dependent regulatory framework illuminates how fundamental differences in chaperone control shape cnidarian stress physiology, offering new mechanistic insight into the cellular basis of coral bleaching susceptibility under ocean warming.

## Introduction

Environmental stress responses vary widely among organisms, enabling some species to tolerate rapid environmental fluctuations while others cannot (1). Anthropogenic climate change, driven by fossil fuel combustion and habitat alteration, is amplifying major environmental stressors such as ocean warming, acidification, hypoxia, salinity shifts, and pollutant exposure (2–3). These changes are disrupting marine ecosystems, impairing organismal physiology, and threatening the stability of oceanic food webs and ecosystem services (3–5). Among the most vulnerable taxa are scleractinian corals and other cnidarians (e.g., sea anemones, jellyfish) whose symbioses underpin the biodiversity and productivity of coral reef ecosystems (6). Rising sea temperatures trigger the breakdown of symbiotic associations through the expulsion of the photosynthetic symbionts, leading to widespread bleaching and mortality (7).

Susceptibility to bleaching varies among symbiotic cnidarians. A central question in marine biology is how cnidarians and their symbiotic partners respond at the molecular level to thermal stress. One of the most conserved cellular defense systems involves heat shock proteins (HSPs), a family of ATP-dependent molecular chaperones that maintain proteostasis by refolding denatured proteins and preventing aggregation (8–9). Discovered by Ferruccio Ritossa in *Drosophila melanogaster* in the 1960s (10–11), HSPs and the associated heat shock response represent one of the most ancient stress response pathways across all domains of life (12–13). This conservation reflects strong evolutionary pressure to maintain cellular homeostasis under stress, with lineage-specific adaptations arising to meet distinct environmental challenges (13). Yet, despite this, the diversity and regulation of HSP expression in cnidarians remain poorly resolved, and we still lack a clear understanding of how the canonical heat shock response has been adapted to the distinct thermal niches and symbiotic lifestyles of these organisms.

Among the known HSP families, the major families (HSP60, HSP70 and HSP90, named according to their molecular weight) play a specialized role in stress mitigation. The canonical HSP90 acts as a constitutive stabilizer of signaling pathways, HSP70 is rapidly and highly inducible during acute stress, and HSP60 functions within mitochondria to refold imported or misfolded proteins under prolonged stress (14–18). In particular, HSP60 is the eukaryotic homolog of the bacterial chaperonin GroEL, suggesting evolutionary roots in prokaryotic proteostasis systems (15–16). Its structure and function are remarkably conserved between fruit fly, chicken, *E. coli,* and yeast (13–16), supporting a role as essential components of mitochondrial homeostasis (17–18). While much is known about the conservation of this protein in particular in mammalian systems, much less is known about how this ancient chaperonin is regulated across evolutionarily distant yet functionally comparable lineages, especially in early-divergent animals such as cnidarians, where mitochondrial stress responses remain poorly characterized.

In marine invertebrates, HSP expression patterns are increasingly used as biomarkers of stress and resilience (19–21). In stony corals, HSP expression and their corresponding encoding genes have been studied in relation to acclimation to elevated temperatures and carbon dioxide levels, while in anemones and jellyfish, HSP dynamics have provided insights into symbiotic flexibility and adaptive physiology (22–25). However, comparative analyses that connect evolutionary conservation of HSP structure with the diversification of its regulatory behavior remain scarce. This disconnect limits our understanding of how conserved molecular components can yield divergent stress phenotypes across lineages.

Here, we first investigated the heat-responsive regulation of mitochondrial heat shock protein 60 (HSP60) across three ecologically and phylogenetically distinct symbiotic cnidarians: the scleractinian coral *Pocillopora acuta*; the model sea anemone *Exaiptasia diaphana*; and the upside-down jellyfish *Cassiopea xamachana*. These species differ in habitat, thermal tolerance, and the nature of their association with *Symbiodiniaceae* symbionts: *P. acuta* forms obligate mutualisms in reef environments, *E. diaphana* maintains facultative symbiosis, and *C. xamachana* perpetuates obligate symbiosis while inhabiting thermally variable mangroves and lagoons (26–29). By comparing their HSP60 expression under standardized heat-stress regimes, we aimed to characterize the conditions of chaperone regulation. Observed cross-reactivity of a mammalian HSP60 antibody across cnidarian taxa, then motivated complementary phylogenetic analyses to assess the evolutionary conservation of this mitochondrial chaperone. Together, this integrative framework links physiological and evolutionary perspectives to explore how a deeply conserved molecular system operates across diverse cnidarian lineages.

## Results

### Organism level HSP60 protein expression across three cnidarians

HSP60 induction was investigated across marine invertebrates belonging to three cnidarian classes (Fig.1). Immunoblotting of whole-tissue lysates detected HSP60 across all three species (β-actin as protein loading control, protein band densitometry normalized to loading control) (Fig. 2). In *P. acuta*, HSP60 was not detected at either the control (25 °C) or heated (30 °C) conditions (Fig. 2A). In *E. diaphana*, transient HSP60 expression was detected under control conditions (22 °C) but gradually increased at 12 and 24 h under heat stress (27 °C; Fig. 2B). In *C. xamachana*, HSP60 expression was constitutively detected across all time points under both ambient (27 °C) and heat stress (32 °C) conditions (Fig. 2C), and with expression gradually increasing with exposure to elevated temperatures. Altogether, these data indicate species-specific basal abundance and heat inducibility of HSP60, with *P. acuta* showing no response with heat stress of +5 °C above its thermal optimum, *E. diaphana* showing temperature-responsive upregulation and *C. xamachana* exhibiting stable expression over the 24 h time course under control conditions and a slight temperature-dependency under heated conditions.

**Figure 1.**
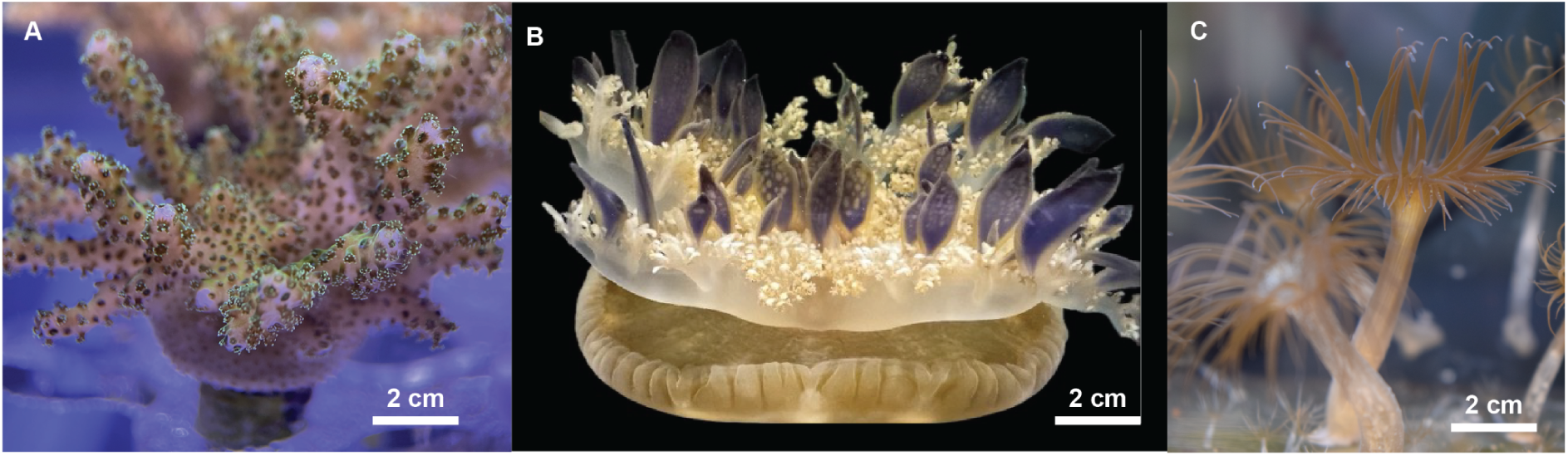
Photographs of symbiotic cnidarians used in this study. (A) *Pocillopora acuta* polyps (Scleractinia), a reef-building coral widely distributed across the Indo-Pacific. (B) *Cassiopea xamachana* medusa (Scyphozoa), upside-down jellyfish are native to the Caribbean and tropical western Atlantic. (C) *Exaiptasia diaphana* (Actiniaria), also known as the glass anemone, is predominantly found on warm-temperate to tropical shores of the Atlantic, Indo-Pacific, and Mediterranean.

**Figure 2.**
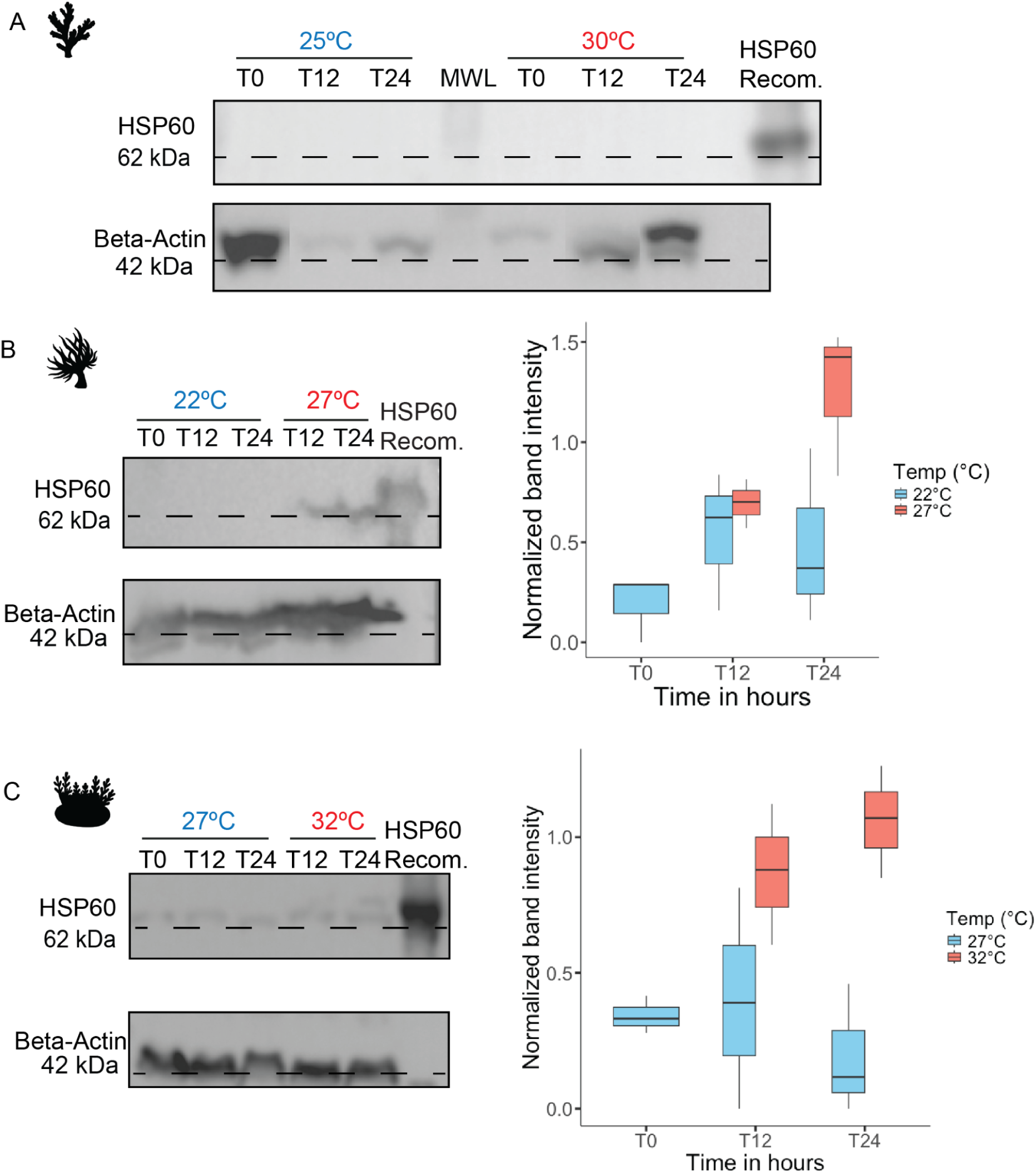
Quantitative Western Blot analysis of HSP60 across three species. Immunoblot detection of HSP60 (∼62 kDa) in tissue extracts at control (blue) and warm (red) temperatures and protein band density graphs (n=3, ± SE) for *P. acuta* (A), *E. diaphana* (B), and *C. xamachana* (C). Samples were taken at various time points during 24 h exposure (T0, T12, T24). Recombinant HSP60 standard (Recom.) was included, and β-actin (∼42 kDa) was used as a loading control.

### HSP60 expression under in vivo and in vitro heat stress in Pocillopora acuta

In contrast to the data presented above, *in vitro* assays of isolated *P. acuta* cells revealed variable HSP60 expression over exposure duration across replicates (Fig. S1) when exposed to heat stress of +5 °C above its thermal optimum (30 °C) with a general temperature-dependent response, and a basal but variable expression under control conditions (25 °C; Fig. 3 and S1). These results indicate a context-dependent difference in HSP60 regulation in *P. acuta*, where isolated cells exhibit a transient, time- and temperature-dependent increase in HSP60 expression under heat stress, whereas intact tissue fragments do not show detectable induction under the same conditions.

**Figure 3.**
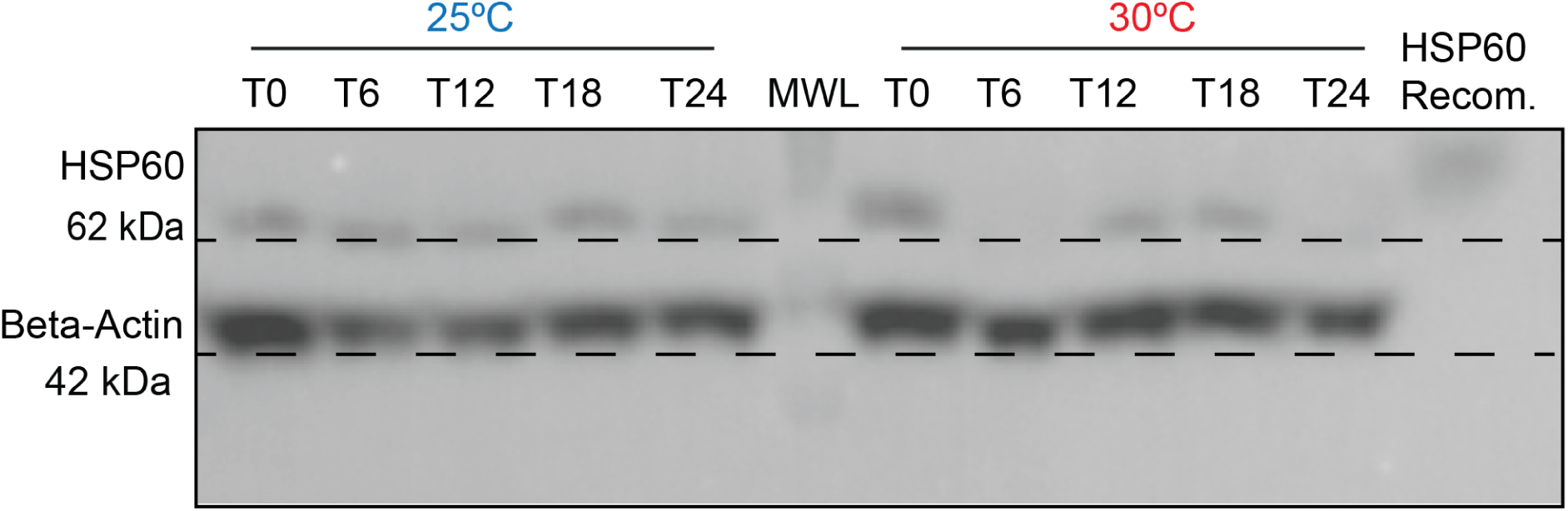
Western Blot analysis of *in vitro* responses of *P. acuta* to acute heat stress. Cellular-level stress assessed by HSP60 immunoblotting under control (blue) and heated (red) conditions and sampled at various time points over 24 h (T0—T24); the HSP60 band is visible at ∼62 kDa, with β-actin (∼42 kDa) as loading control. “Recom.” denotes the recombinant HSP60 standard, and MWL indicates the molecular-weight ladder.

### Multiple Sequence Alignment reveals high conservation of functional motifs

The candidate HSP60 orthologs were identified in *P. acuta* (ID: TCONS_00030188), *E. diaphana* (ID: P18687), and *C. xamachana* (ID: Casxa1|9735) through BLASTp searches against the canonical human HSPD1 sequence (UniProt ID: P10809). All three cnidarian proteins exhibited strong homology to the mitochondrial chaperonin 60 family (query coverage ≥ 80 %), confirming their identity as HSP60 orthologs. Multiple sequence alignment (MSAs) between the cnidarian and human HSP60 proteins revealed extensive conservation across key functional motifs, with over 85% sequence identity across the aligned regions (Fig. 4, S7), indicating high structural and functional similarity. Furthermore, sequence comparison across the full0length protein demonstrated overall conservation ranging from 69.6% to 91% identity and 84% to 96% positives relative to human HSPD1 (Table S7).

**Figure 4.**
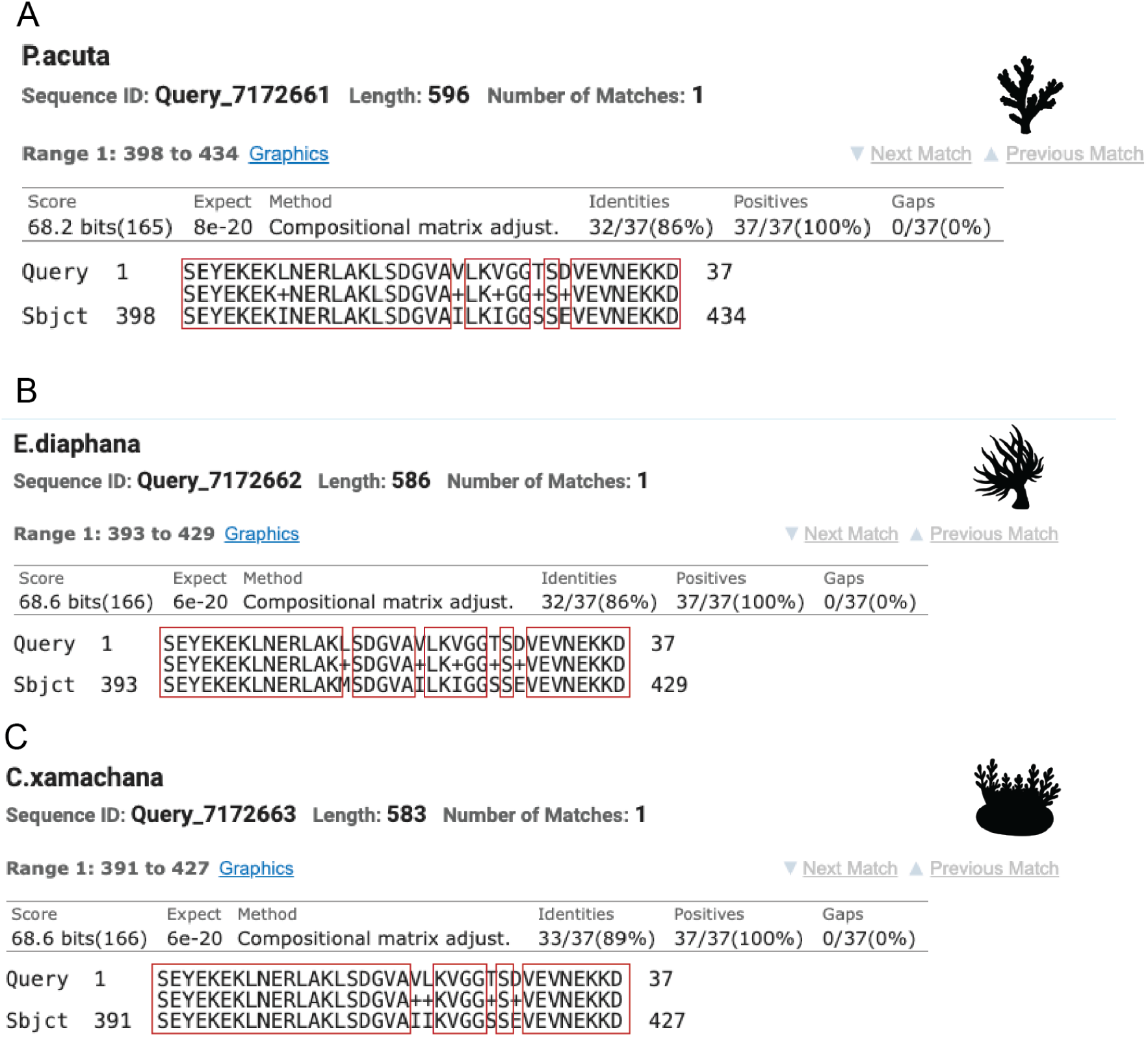
Conserved HSP60 motif shared across Cnidaria and humans. Pairwise BLASTp alignments of human mitochondrial HSP60 (HSPD1) with HSP60 orthologs from (A) *P. acuta* (596 aa), (B) *E. diaphana* (586 aa), and (C) *C. xamachana* (583 aa) reveal a highly conserved 37 amino acid segment within the C-terminal region (alignment ranges: ∼391–434). Identities are 86% (*P. acuta* and *E. diaphana*) and 89% (*C. xamachana*), with 100% positives and no gaps (E-values between 8×10^−20^ and 6×10^−20^).

Epitope analysis of the experimentally validated human HSP60 antibody region (amino acids 383-419, Thermo Scientific) showed partial to complete conservation across the corresponding cnidarian residues (Fig. S4, Table S8). Predicted epitope identity scores (86% for *P. acuta* and *E. diaphana*, 89% for *C. xamachana*) further support strong antibody cross-reactivity potential and underscore the deep conservation of antigenic and functional domains within the HSP60 family.

### A structurally conserved HSP60 epitope spans Cnidaria and Mammalia

To connect sequence conservation to structural context, we predicted HSP60 tertiary structures for *P. acuta*, *E. diaphana*, and *C. xamachana* using AlphaFold3 and mapped the 37-residue antigenic motif identified by pairwise alignments to human HSP60 (HSPD1, Fig. 5) (30). AlphaFold consistently recovered the canonical domain organization across all three cnidarians, placing the conserved epitope in an equivalent position on the equatorial domain surface (Fig. 5). This structural correspondence provides a clear rationale for the observed antibody cross-reactivity and indicates strong evolutionary constraint on this region. The equatorial domain mediates ATP binding and oligomerization in group I chaperonins (31–32), and conservation of its surface-exposed epitope across taxa supports an essential mechanistic role. All predicted cnidarian HSP60 models display the canonical architecture with an extended C-terminal tail (Fig. 5), within which the conserved motif remains solvent exposed on the monomer surface. The shared positioning and accessibility of this epitope among cnidarians and mammalian homologs highlight deep functional conservation of the HSP60 complex across Metazoa.

**Figure 5.**
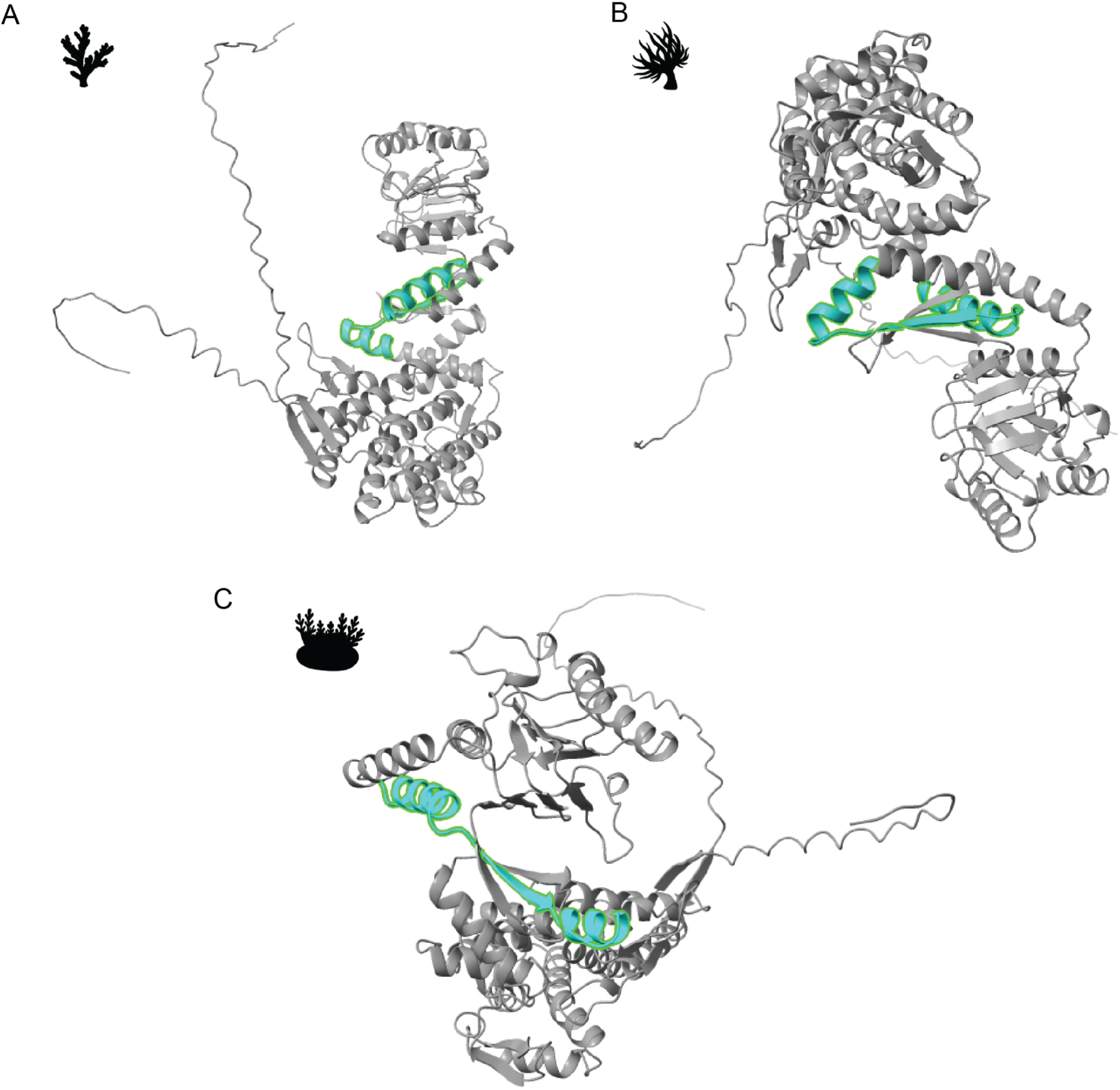
AlphaFold-predicted tertiary structures of HSP60 homologs in three cnidarian species. Monomeric HSP60 models for (A) *P. acuta*, (B) *E. diaphana*, and (C) *C. xamachana* generated using AlphaFold3. The conserved 37-residue epitopes (green) identified by pairwise alignments are highlighted in each model and map to an equivalent region within the equatorial domain. Per-residue confidence (pLDDT) and predicted aligned error (PAE) are consistent with high confidence in the globular cores and modestly lower confidence at hinge/loop regions (pTM =0.73 for *P. acuta*, pTM = 0.76 for *E. diaphana* and pTM = 0.78 for *C. xamachana*). Canonical HSP60 functions *in vivo* as a tetradecameric double ring; only the monomeric fold is shown here for clarity.

### Phylogenetic analysis supports evolutionary conservation across Cnidaria

Maximum-likelihood phylogenetic analysis of aligned HSP60 sequences recovered the expected relationships among the three cnidarians (Fig. S5). The coral *P. acuta* (Scleractinia) and the sea anemone *E. diaphana* (Actiniaria) formed a well-supported sister clade, consistent with the established anthozoa phylogeny (33). The jellyfish *C. xamachana* branched separately, reflecting its placement within the medusozoan lineage. The tree was rooted with the human mitochondrial HSP60 (HSPD1/CH60), which served as an outgroup to Cnidaria. This topology confirms that cnidarian HSP60 proteins are orthologous to the human homolog and that the locus retains sufficient phylogenetic signal to resolve divergence within Cnidaria.

### TargetP-2.0 analysis confirms mitochondrial localization of candidate HSP60 sequences

To verify subcellular targeting of the cnidarian HSP60 candidates, we used TargetP-2.0 prediction tool, which estimates the likelihood of a protein being mitochondrial in origin. Given that HSP60 is a mitochondrial chaperonin in mammals, this analysis was performed to confirm whether the cnidarian homologs share the same localization.

Predictions for *P. acuta* indicate a high mitochondrial targeting probability (0.99) with a predicted cleavage site between amino acids 28-29 (VGF-ST; Fig. S6A). *E. diaphana* showed a similarly strong probability (0.97) with a predicted cleavage site also between its amino acids 28-29 (AXF-ST; Fig. S6B), and *C. xamachana* yielded a 0.99 probability with a cleavage site between residues 26-27 (ASF-ST; Fig. S6C). In all cases, the predicted N-terminal transit peptide and corresponding cleavage positions align with canonical mitochondrial targeting motifs observed in metazoan HSP60s. These consistent, high-confidence predictions (e.g., 0.99 probability for *P.* acuta and *C. xamachana*, 0.97 for *E. diaphana*) support that cnidarian HSP60 sequences encode mitochondrial chaperonins and reinforce the evolutionary conservation of subcellular localization within this protein family.

## Discussion

Our findings demonstrate that mitochondrial HSP60 is deeply conserved across at least three cnidarian classes and maintains clear mechanistic continuity with its mammalian homolog. Using an anti-HSP60 monoclonal antibody (IgG mouse clone LK-2, Thermo Scientific) designed for mammalian systems, we successfully detected HSP60 in three ecologically and phylogenetically distinct cnidarians—*Pocillopora acuta*, *Exaiptasia diaphana*, and *Cassiopea xamachana—*establishing functional cross-reactivity and supporting deep evolutionary conservation (Fig. 2-3). Pairwise sequence alignments identified a 37-residue motif with high identity to human HSP60 (∼86–89%) and even greater conservation among cnidarians (∼92–95%). (Fig. 4 and S4). AlphaFold3-based structural mapping localized this motif to the equatorial domain in an equivalent position across taxa, consistent with antibody recognition and strong structural constraint (Fig. 5). These data confirm that HSP60’s core mitochondrial chaperon function, in partnership with HSP10, is conserved from cnidarians to mammals, in agreement with previous observation in metazoan and bacterial studies (34–39). This conservation of the domain, which mediates ATP binding and oligomerization in Group I chaperonins, further provides strong evidence that the co-chaperonon HSP10 exists and functions equivalently in these cnidarians, consistent with deep mechanistic continuity across Metazoa. Conservation of this domain implies that secondary roles of HSP60, such as those involved in apoptosis, immune modulation, and unfolded-protein responses, likely originated early in animal evolution (40–44).

Across experiment systems, HSP60 regulation under heat stress was context-dependent rather than uniform. We applied a +5 °C increase above each species’ laboratory growth temperature to standardize stress intensity. Under *in vivo* conditions, *P. acuta* did not display detectable HSP60 expression, *E. diaphana* showed gradual HSP60 accumulation over time with exposure to heat stress, whereas *C. xamachana* maintained constitutive levels across treatments with a temperature-dependency. In contrast, *in vitro* exposure of *P. acuta* cells resulted in a variable HSP60 expression under both control and heat-stressed conditions (5 °C above thermal optimum). These differences are best explained by biological rather than technical factors. In dissociated cells, tissue-level signaling, metabolic coupling, and intercellular feedback are disrupted, leading to altered stress perception and loss of physiological buffering (45–47). Consequently, equivalent thermal challenges may trigger adaptive HSP60 induction in isolated cells compared to intact tissue. Comparable phenomena are observed in mammalian systems, where cell detachment affects HSF1 activation and alters HSP60 and HSP70 induction due to disrupted integrin signaling and mitochondrial instability (48–51). This cross-system parallel suggests that HSP60 regulation is fundamentally dependent on integrated physiological context, tissue organization, and taxa rather than temperature alone, supporting a hierarchical model of stress regulation that is altogether context-, lineage-, and temperature-dependent. For this thermally sensitive *P. acuta*, the absence of detectable HSP60 induction in intact tissue fragments suggests that physiological buffering or successful tissue integrity signaling suppresses the immediate initiation of mitochondrial stress and secondary stress pathways (including those involving HSP60’s roles in apoptosis and immune modulation). This suppression, however, prevents the deployment of a critical molecular defense mechanism necessary to manage proteostatic demand, thereby contributing to the species’ thermal sensitivity and known susceptibility to bleaching

The sea anemone *Exaiptasia* spp. (previously *Aiptasia*) exemplifies this hierarchical organization of stress regulation across environmental contexts and symbiotic states. Previous studies show that *E. diaphana* engages canonical signaling pathways such as TGF-β, NF-κB, and p53 during temperature- and salinity-driven stress (52–54). Under thermal extremes, this species exhibits cell-type- and symbiotic-stage–dependent transitions from apoptosis to necrosis, accompanied by HSP upregulation that likely mitigates programmed cell death (55). Environmental salinity further modulates these trajectories: while hypersalinity is generally deleterious in corals (often reducing photosynthetic efficiency and inducing bleaching) (56–58) moderate-to-high salinity in *Exaiptasia* reduces symbiont loss and heat-induced bleaching, correlating with microbiome shifts that enhance nitrogen retention and antioxidant capacity (59). During prolonged heat exposures (e.g., 34 °C for 28 days), stress pathways diverge by symbiotic state, indicating that oxidative stress is jointly governed by symbiotic state and organismal integration level (54).

Jellyfish of the genus *Cassiopea* provide a complementary perspective on thermal adaptation. Living in shallow mangrove habitats characterized by strong diel temperature fluctuations, *C. xamachana* is physiologically pre-conditioned to thermal variability. Previous work reports minimal differential HSP expression under acute thermal stress and enhanced thermotolerance following acclimation to ∼32 °C (60). Similarly, *Cassiopea andromeda* maintains symbiosis during long-term exposure to ∼ +6 °C, with high chlorophyll *a* content in symbionts indicating sustained photosynthetic function (61). Nevertheless, bleaching can occur under prolonged or extreme regimes: *C. xamachana* polyps bleach at 35 °C for two weeks, and adults at 34 °C for one week exhibit reduced bell diameters and partial bleaching, although reports of *Cassiopea* bleaching in the field remain rare (62–64). Comparisons with other scyphozoans, such as *Aurelia labiata* (ephyrae), which exhibit growth and swimming impairments at only +1.5 °C above optimum, underscore the exceptional thermotolerance of *Cassiopea* spp (65). The constitutive and temperature-dependent HSP60 signal observed in *C. xamachana* aligns with its life-history-driven thermal resilience.

Together, our results reveal a unifying pattern: HSP60’s amino acid sequence and domain architecture are evolutionarily conserved from cnidarians to mammals, but its regulatory dynamics are modulated by lineage, stress regime (intensity, duration, modality), and the level of biological organization. This context dependence is biologically meaningful because HSP60 abundance reflects mitochondrial integrity, free radical balance, and proteostatic demand (66). The transient HSP60 signal found in heated *P. acuta* cells likely reflects fluctuations in mitochondrial import efficiency or membrane potential with heat stress onset (67), while the absence of signal in *P. acuta* fragments under heat stress confirms a context-dependent suppression mechanism that contributes directly to the species’ thermal sensitivity and high susceptibility to bleaching (68). Thus, HSP60 functions not only as an evolutionarily ancient chaperone but also as a sensitive biomarker for mitochondrial stress and biological organization.

Importantly, our findings highlight the interpretive value of negative and variable results. While strong HSP induction under severe stress is well documented in scleractinian corals, few studies explicitly report conditions under which HSP induction fails (69–72). Documenting cases of non-induction, such as in *P. acuta* fragments and dissociated cells exposed to +5 °C above optimum (variation in expression over exposure time), is essential for defining boundaries on HSP60’s reliability as a stress biomarker and improving reproducibility in coral stress research (73).

The future work should integrate protein- and transcript-level measurements (e.g., qPCR, RNA-seq), employ targeted proteomics for absolute quantification, and localize HSP60 via immunofluorescence using mitochondrial markers and peptide-blocking controls. Physiological assays for free radical generation, mitochondrial membrane potential, ATP/ADP ratio, and intracellular pH, combined with apoptosis markers (caspase activity, TUNEL), would clarify the mechanistic basis of HSP60 regulation. Implementing ecologically realistic stress paradigms, such as gradual temperature ramps, fluctuating salinity and pH, and altered photoperiod, under defined symbiotic states, coupled with comparative genomics (orthology validation, genome-wide dN/dS and site-specific constraint analyses) will help delineate when HSP60 reliably indicates mitochondrial stress versus regulatory fine-tuning. Such integrative approaches will refine the application of HSP60 as a mechanistic biomarker for predicting symbiosis stability and bleaching susceptibility under ocean warming.

## Material and Methods

All product details can be found in the Supplementary Materials associated with each methodological section.

### Study organisms overview

The organisms used in this study are the scleractinian coral *Pocillopora acuta* (green phenotype), sea anemone *Exaiptasia diaphana* (H2 clone line), and the upside-down jellyfish *Cassiopea xamachana* (Fig. 1). *P. acuta* is considered to be a thermally sensitive species, with documented bleaching responses at temperatures 1-2 °C above local summer maxima (74–75). The symbiont population in *P. acuta* is typically composed of *Cladocopium goreaui* (C1), with a low relative abundance of *Durusdinium* type D1 (76–79). The colonies used in this study belong to a monoclonal population originating from Hawaii and propagated in aquaculture for research purposes (Putnam Lab, University of Rhode Island, USA).

The glass anemone *E. diaphana* is a small, pale brown, symbiotic anemone, which has been widely used as a model organism for understanding cnidarian symbiosis and physiology (80). This species is commonly associated with the dinoflagellate *Breviolum minutum* (formerly Clade B), but is capable of hosting other *Symbiodiniaceae* genera under experimental conditions, e.g, *Symbiodinium* spp., *Durusdinium* spp. (81–82). H2 clone line individuals were originally shared with us from the Jinkerson Lab (University of California, Riverside, USA).

The upside-down jellyfish *C. xamachana* is a rising model system for investigating cnidarian-algal symbiosis, tissue regeneration, and evolutionary biology. Recent investigations highlight the use of *Cassiopea* spp. organisms as indicator species for environmental monitoring and ecotoxicology, and virology (83–87). *C. xamachana* is mostly found in the Western Atlantic Ocean, Caribbean Sea, Gulf of Mexico, and Florida Keys (specimens used here), and hosts *Symbiodinium microadriaticum* (clade A1). *S. microadriaticum* infection is essential to *Cassiopea* spp. life cycle (88). Polyps of *C. xamachana* were originally shared with us from the Buckley Lab (Auburn University, USA).

### Experimental design

Two complementary approaches were used to assess HSP60 regulation: *in* vitro assays using dissociated cells from *P. acuta* to capture cellular-level responses, and *in* vivo, whole-organism experiments from which tissue slurries were prepared using *P. acuta, C. xamachana,* and *E. diaphana*.

*In vitro* assays were performed by exposing dissociated cells to control and heated conditions (+5 °C above ambient aquaria temperature) in lit cell culture incubators (12 h-12 h photoperiod). *In vivo* experiments were conducted by transferring the whole organisms from their ambient aquaria directly to a heated experimental tank, with a temperature of +5 °C above ambient aquaria temperature: *P. acuta* control = 25 ±0.5 °C vs. heated = 30 ±0.5 °C, *E. diaphana* control = 22 ±0.5 °C vs heated 27 ±0.5 °C, and *C. xamachana* control 27 ±0.5 °C vs. heated = 32 ±0.5 °C. Dissociated cells, tissue slurries and lysates were obtained as described next.

### Coral cell dissociation

The coral cell dissociation protocol is detailed in Supporting information (S3) and adapted from Roger *et al.* (89). In brief, a small piece (or nubbin) of *P. acuta* fragment was rinsed with sterile-filtered artificial seawater (FASW) and incubated in Ca-Mg-free FASW (CMF-FASW) for 1 h. Post incubation, the nubbin was gently washed with CMF-FASW to detach the cells from the surface of the skeleton. The resulting cell suspension was then centrifuged and resuspended in coral cell culture media before being transferred to a clear multi-well plate for incubation under experimental conditions. Once the experiment was completed, the cell suspensions were centrifuged at 8000 rpm (6010 RCF) for 10 minutes at 4 °C. The supernatant was removed, and the cell pellets were stored in -20 °C until protein extraction.

### Tissue homogenization

Once the thermal experiment was completed, individuals of *E. diaphana* (∼5 mm) and *C. xamachana* (∼2 cm in diameter) were transferred to a 50 mL conical tube containing 4 mL FASW and homogenized (handheld motor) for 10-15 second intervals on ice to prevent protein degradation. The homogenate was centrifuged at 8000 rpm (6010 RCF) for 10 minutes at 4 °C. The supernatant was removed, and the cell pellets were stored in -20 °C until protein extraction.

### Total protein extraction and analysis

The total protein extraction for dissociated cells from the *P. acuta* fragment was adapted from Seveso *et al.* (69), with minor modifications. The cells were resuspended in 100 µL of lysis buffer (0.0625 M Tris-HCl, pH 6.8, 10 % glycerol, 0.1% w/v SDS, 1% w/v 2-mercaptoethanol containing 1 mM phenylmethylsulfonyl fluoride, and complete EDTA-free cocktail of protease inhibitors). The cells with the lysis buffer mixture were boiled for 10 minutes at 95 °C and centrifuged at 13,000 rpm (13,523 RCF) for 15 minutes at 4 °C. The supernatant was further clarified by centrifuging again (5,000 rpm or 2348 RCF for 5 minutes).

The total protein concentration was determined using the Bradford Assay, and the absorbance was measured using a microplate reader (595 nm).

The protein samples were separated by sodium dodecyl sulfate-polyacrylamide gel electrophoresis (SDS-PAGE), using 10% polyacrylamide gels; 20 µg of total protein was loaded into each lane of the gel. Two gels were run simultaneously; one gel was stained with Instant Blue Coomassie Protein Stain to visualize protein bands, and the other gel was used for the western immunoblot assay. The PVDF membrane (0.2 µm) was stained with Ponceau S staining solution to confirm protein transfer on the membrane. The membrane was incubated in 5% Blocking solution (5 % nonfat dry milk in 1X TBST: Tris-Buffered Saline, 0.1 % Tween-20) for 1 h. Following blocking, the membrane was incubated with Anti-HSP60 monoclonal antibody (IgG mouse clone LK-2) at a 1:1000 dilution, and the membrane was triple-washed with 1X TBST buffer and incubated with horseradish peroxidase conjugated-anti-mouse IgG secondary antibody, 1:10,000 dilution in 5% blocking solution for 2 hours. The chemiluminescence signal was detected using Pierce ECL western blotting substrate and imaged using a gel imager. The membranes were stripped, reblocked, and incubated with anti-actin monoclonal antibody (clone C4) at a 1:3000 dilution overnight at 4°C. The membrane was triple-washed with 1X TBST buffer and incubated with horseradish peroxidase conjugated-anti-mouse IgG secondary antibody, 1:10,000 dilution in 5% blocking solution for 2 h. The chemiluminescence signal was detected using Pierce ECL western blotting substrate and imaged using a gel imager.

### Bioinformatics

#### Multiple Sequence Alignment

To assess evolutionary conservation and sequence homology of HSP60 across representative cnidarian taxa, multiple sequence alignment (MSA) using pairwise alignment was performed using HSP60 amino acid sequences derived from *P. acuta*, *E. diaphana*, and *C. xamachana*, alongside the canonical human HSP60 (HSPD1, UniProt ID: P10809) as reference. The candidate coral (*P. acuta*) sequences were obtained from Vidal-Dupiol *et al.* (90), *C. xamachana* sequences from the Joint Genome Institute (https://jgi.doe.gov/), and *E. diaphana* sequences from ReefGenomics (91–92). Protein identity and orthology were confirmed via BLASTp (NCBI) against the non-redundant database, retaining top hits with query coverage >80 % and e-value <1e^-5^. Conserved residues and functional domains were annotated based on the human HSP60 reference sequence mentioned above.

#### Epitope alignment

For antibody epitope mapping, the experimentally validated linear epitope of human HSP60 (mouse, monoclonal LK-2, MA5-45114, Thermo Scientific) was obtained from product datasheets (Thermo Scientific). The corresponding epitope region (amino acid 383–419) was aligned to the cnidarian HSP60 orthologs within the MSA framework to identify conserved and variable residues potentially contributing to antibody recognition. Phylogenetic relationships among cnidarian HSP60 sequences and the human HSPD1 reference were to be inferred using MEGA X (93) under the Maximum Likelihood (ML) method with default parameters. Branch support was assessed using 1,000 bootstrap replicates.

#### AlphaFold3 structural prediction and epitope mapping

Predicted tertiary structures of HSP60 homologs from *P. acuta, C. xamachana,* and *E. diaphana* were obtained from the AlphaFold Protein Structure database (DeepMind and EMBL). For each species, the highest-ranked model was downloaded (AlphaFold Version 4) in CIF format and visualized in UCSF ChimeraX (Version 1.10.1). Per-residue confidence scores were derived from the predicted local distance difference test (pLDDT) values.

Conserved regions corresponding to the human HSP60 epitope (amino acid 383-419) were identified using Clustal Omega multiple sequence alignment (90). Homologous residues were mapped to each cnidarian HSP60 sequence (positions 398–434 in *P. acuta*, 393–429 in *E. diaphana,* and 391–427 in *C. xamachana*) and highlighted on their respective AlphaFold-predicted tertiary structures. These residues were rendered in cartoon representations (cyan) to evaluate spatial conservation, solvent exposure, and structural context of the conserved epitope within the chaperonin architecture.

#### Mitochondrial protein prediction

To determine the subcellular localization of cnidarian HSO60 homologs, candidate protein sequences were analyzed using TargetP-2.0 (DTU Health Tech (https://services.healthtech.dtu.dk/services/TargetP-2.0/). Sequences from *P. acuta* (TCONS_003018), *E. diaphana* (P18687), and *C. xamachana* (Casxa1|9735) were submitted as the queries to predict mitochondrial targeting peptides. TargetP-2.0 provides the probability of mitochondrial localization and identifies N-terminal cleavage sites. Predicted probability scores and cleavage positions were recorded for each sequence and visualized as probability plots of amino acid position versus localization likelihood (Fig. S6).

## Supplementary materials

**S1** Western blot quantification of protein abundance in cultured *P. acuta* cells across a 24-h temperature treatment.

**S2** Animal husbandry.

**S3** Coral cell dissociation and cell sample collection.

**S4** Conservation of C-terminus HSP60 antigenic epitope across study species.

**S5** Phylogenetic placement of cnidarian HSP60 proteins.

**S6** TargetP-2.0 outputs confirming mitochondrial localization.

**S7** Table showing alignment of candidate sequences.

**S8** Table showing epitope alignments among candidate sequences.

## Data availability

The protein sequences for the study organisms were obtained from Reef Genomics (http://reefgenomics.org/), Joint Genome Institute (https://jgi.doe.gov/), and NCBI (https://www.ncbi.nlm.nih.gov/). The detailed protocol for S3 is available on Protocols.io. (https://www.protocols.io/private/1AB2BFF1ADF211F0AA040A58A9FEAC02)

The immunoblot quantification data and summary of experiments are available on Open Science Forum (https://osf.io/zwp7g/files).

All analyses were performed using R statistical software (Version 2025.09.1+401 (2025.09.1+401). The western blot quantification was performed on ImageJ.

## Acknowledgments

We thank Dr. Hollie Putnam, Dr. Robert Jinkerson, and Dr. Megan Maloney for providing animal cultures and husbandry techniques. We thank our graduate research volunteer, Charitha Rajapakse, for assisting with total protein analysis. We thank all the graduate and undergraduate student volunteers of ASU’s Marine Biochem Research Lab (Roger Lab) for assisting in the maintenance of the aquaria and organisms.

