## Supplementary Information for "Deep conservation of mitochondrial HSP60 structure with lineage-specific and context-dependent regulation reflects thermal resilience in cnidarians"

**List of supporting information:**

**S1.** Western blot quantification of protein abundance in cultured *P. acuta* cells across a 24 h temperature treatment.

**S2.** Animal husbandry protocols.

**S3.** Coral cell dissociation and cell sample collection.

**S4.** Conservation of C-terminus HSP60 antigenic epitope across study species.

**S5.** Phylogenetic placement of cnidarian HSP60 proteins.

**S6.** TargetP-2.0 outputs confirming mitochondrial localization.

**S7.** Table showing alignment of candidate sequences.

**S8.** Table showing epitope alignments among candidate sequences.

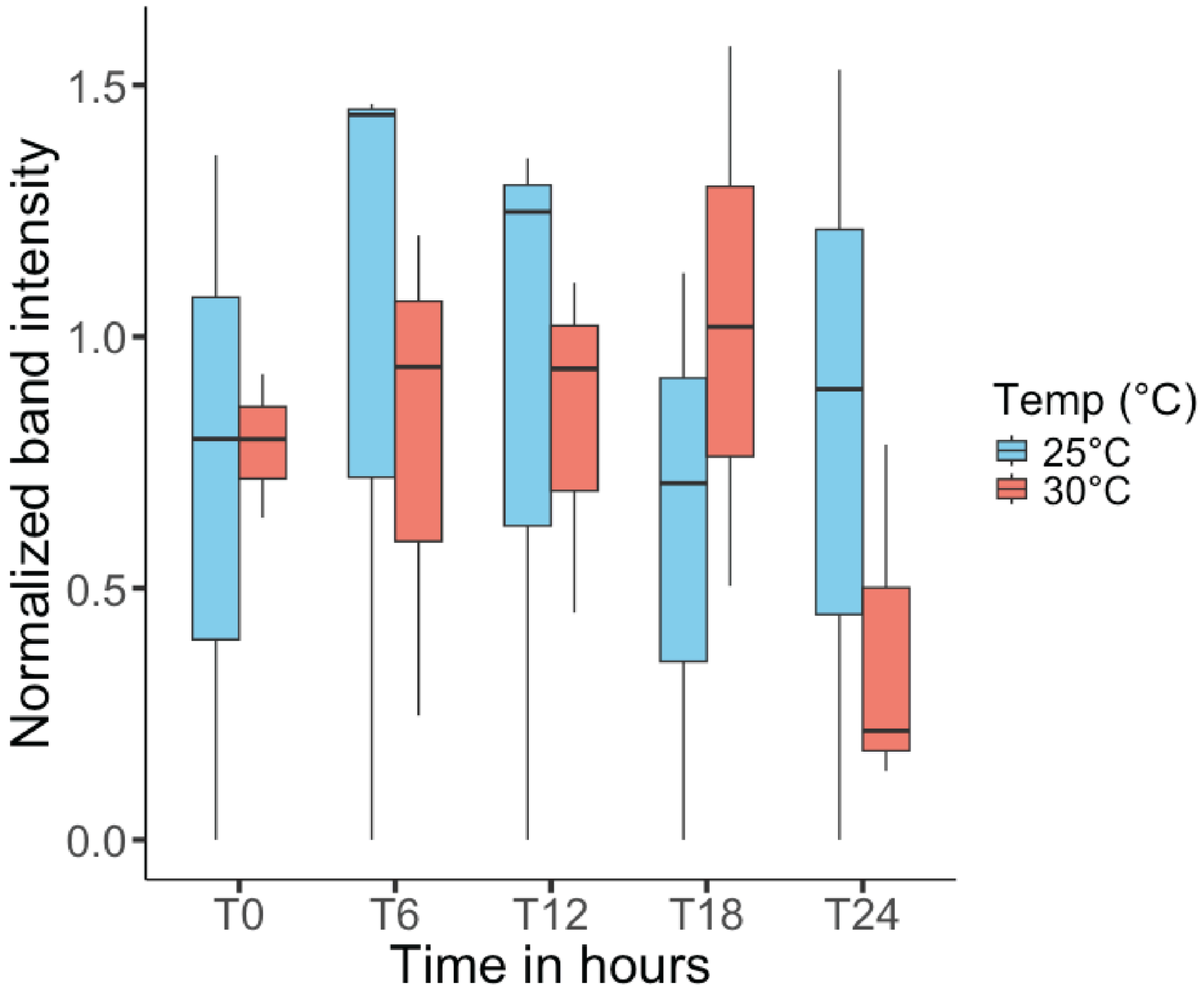


**Figure S1. Western blot quantification of protein abundance in cultured *P. acuta* cells across a 24-h temperature treatment.** Cells were maintained at 25 °C (blue) or 30 °C (red) and sampled at 0, 6, 12, 18, and 24 h (T0–T24). Band intensities were quantified by densitometry, normalized to the loading control (β-actin), and plotted as normalized intensity. (n=3, ±SE)

**S2. Animal Husbandry**

Colonies of *P. acuta* (green phenotype) were kept in 170 L aquaria (with 24 L sump), supplied with artificial seawater (ASW, Tropic Marin Pro-Reef Salt, Germany, 35 ppt salinity), maintained at 25 ºC (600 W titanium aquarium heater with Inkbird ITC–306 A InkBird temperature controller, USA), under a 12 h: 12 h photoperiod at 150-200 µmol photons $m^{-2}$ $s^{-1}$ irradiance (AP9x LED lights, Kessil, USA). The ASW was constantly filtered (200 µm sock filters, Aquatic Experts), skimmed for organics (Octo Classic 110S protein skimmer, Reef Octopus, USA), and UV sterilized (Aqua Ultraviolet, USA). Seawater parameters (pH, redox potential, temperature, carbonate hardness (kH), calcium, magnesium, phosphate, and nitrate levels) were monitored daily (Proflux 4 GHL, Germany, and API saltwater test kits, USA) and adjusted using dosing buffers when needed (Fauna Marin Balling Light Set, Fauna Marin, Germany). Corals were fed once a week with live *Artemia salina* nauplii (E-Z EGG, Brine Shrimp Direct, USA). Aquaria were cleaned once a week to limit biofouling.  A 10% water change was also done weekly. The colonies used in this study belong to a monoclonal population originating from Hawaii and propagated in aquaculture for research purposes (Putnam Lab, University of Rhode Island).

Specimens of *E. diaphana* (H2 clone line) were maintained in Bisphenol-A-free (BPA-free) clear polypropylene containers (Cambro, USA) supplied with ASW (salinity of 35 ppt, Tropic Marine Pro-Reef salt, Germany), maintained at 22 ºC, and irradiance of 200 µmol photons $m^{-2}$ $s^{-1}$ on a 12 h:12 h photoperiod (Fluval Marine Nano LED light, Germany). The anemones were fed with live *Artemia salina* napulii once a week. The containers were cleaned, and the ASW was renewed with a weekly 100% water change to remove debris, prevent algal growth, and microbial contamination, thereby maintaining optimal conditions for growth and survival of the organisms.

Stock cultures of medusae of *C. xamachana* (non-clonal) were maintained in 170 L aquaria (with 24 L sump) supplied with ASW (Tropic Marin Pro-Reef Salt, Germany, 38 ppt salinity), maintained at 27 ºC (600 W titanium aquarium heater with Inkbird ITC–306 A InkBird temperature controller, USA), under a 12 h: 12 h photoperiod at 150-200 µmol photons $m^{-2}$ $s^{-1}$ irradiance (AP9x LED lights, Kessil, USA). The ASW was constantly filtered (200 µm sock filter, Aquarium Experts), skimmed for organics (OCTO classic 110 SSS protein skimmer, Reef Octopus, USA), and UV sterilized (Aqua Ultraviolet, USA). The seawater parameters (pH, redox potential, temperature, carbonate hardness, calcium, magnesium, phosphate, and nitrate levels) were monitored daily (Profilux 4, GHL, Germany, and API saltwater test kits, USA) and adjusted using buffers when needed (Fauna Marin Balling Light Set, Fauna Marin, Germany). The polyps of *C. xamachana* were grown in a separate container (BPA-free clear Cambro), under the same conditions described above. The polyps were fed with live *Artemia salina* nauplii (E-Z EGG, Brine Shrimp Direct, USA) twice a week, and the medusae were fed daily.

**S3. Coral Cell dissociation and cell sample collection**

(DOI: https://www.protocols.io/private/1AB2BFF1ADF211F0AA040A58A9FEAC02)

A single nubbin from the *P. acuta* colony (~1 cm length) was cut using sterile clippers and transferred to a crystallization dish containing ASW with ReefDip Coral disinfectant (25 µL per mL of ASW) under constant bubbling for 10 minutes. The nubbin was then rinsed with sterile-filtered artificial seawater (FASW) (0.22 µm) thrice and incubated in sterile-filtered calcium-magnesium-free artificial seawater (CMFASW, 0.22 µm) for 1 h in a biosafety cabinet under ambient light. Post incubation, the nubbin was gently washed with CMFASW to detach cells from the surface of the nubbin. The cell suspension was centrifuged at 1200 rpm (204 RCF) for 3 minutes at 25 ºC. The supernatant was removed and the cells resuspended in complete coral cell culture media (15% Dulbecco's Modified Eagle Medium (DMEM) without phenol red, Thermo Scientific Catalog no. 21063029, 10% Fetal Bovine Serum (FBS), ThermoFisher Scientific, Catalog no. A5670701, 0.5%, Antibiotics-antimycotics, Thermo Scientific, Catalog no. 15240062, 0.5% Gentamicin, Thermo Scientific, Catalog no. 15710064, and 74% of sterile FASW.) The coral cell suspension was collected in microcentrifuge tubes every 6 h for a total duration of 24 h. The cells were centrifuged at 8000 rpm (6010 RCF) at 4 ºC for 10 mins. The anemone and jellyfish sample specimens were collected every 12 h, homogenized, and centrifuged at 8000rpm (6010 RCF) at 4 ºC for 10 minutes to collect the tissue pellets, which were maintained at –20 ºC until further use.

A

**E.diaphana_Epitope**

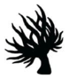

Sequence ID: **Query_5089354** Length: **37** Number of Matches: **1**

Range 1: 1 to 37 Graphics ▼ Next Match ▲ Previous Match

| Score | Expect | Method | Identities | Positives | Gaps |
|---|---|---|---|---|---|
| 68.9 bits(167) | 8e-24 | Compositional matrix adjust. | 35/37(95%) | 37/37(100%) | 0/37(0%) |

```
Query  1    SEYEKEKINERLAKLSDGVAILKIGGSSEVEVNEKKD  37
            SEYEKEK+NERLAK+SDGVAILKIGGSSEVEVNEKKD
Sbjct  1    SEYEKEKLNERLAKMSDGVAILKIGGSSEVEVNEKKD  37
```

B

**C.xamachana_Epitope**

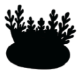

Sequence ID: **Query_5089355** Length: **37** Number of Matches: **1**

Range 1: 1 to 37 Graphics ▼ Next Match ▲ Previous Match

| Score | Expect | Method | Identities | Positives | Gaps |
|---|---|---|---|---|---|
| 68.2 bits(165) | 2e-23 | Compositional matrix adjust. | 34/37(92%) | 37/37(100%) | 0/37(0%) |

```
Query  1    SEYEKEKINERLAKLSDGVAILKIGGSSEVEVNEKKD  37
            SEYEKEK+NERLAKLSDGVAI+K+GGSSEVEVNEKKD
Sbjct  1    SEYEKEKLNERLAKLSDGVAIIKVGGSSEVEVNEKKD  37
```

**Figure S4. Conservation of C-terminus HSP60 antigenic epitope across study species.** Pairwise BLASTp alignments using a 37 amino acid epitope from *P. acuta* HSP60 (query) show strong conservation in (A) *E. diaphana* (35/37 identities; 95% identity; 100% positives; 0 gaps; E-value $8\times10^{-24}$) and (B) *C. xamachana* (34/37 identities; 92% identity; 100% positives; 0 gaps; E-value $2\times10^{-23}$). The high sequence identity across Anthozoa and Medusozoa supports cross-reactivity of anti-HSP60 antibodies and highlights broad phylogenetic conservation of this epitope among cnidarians.

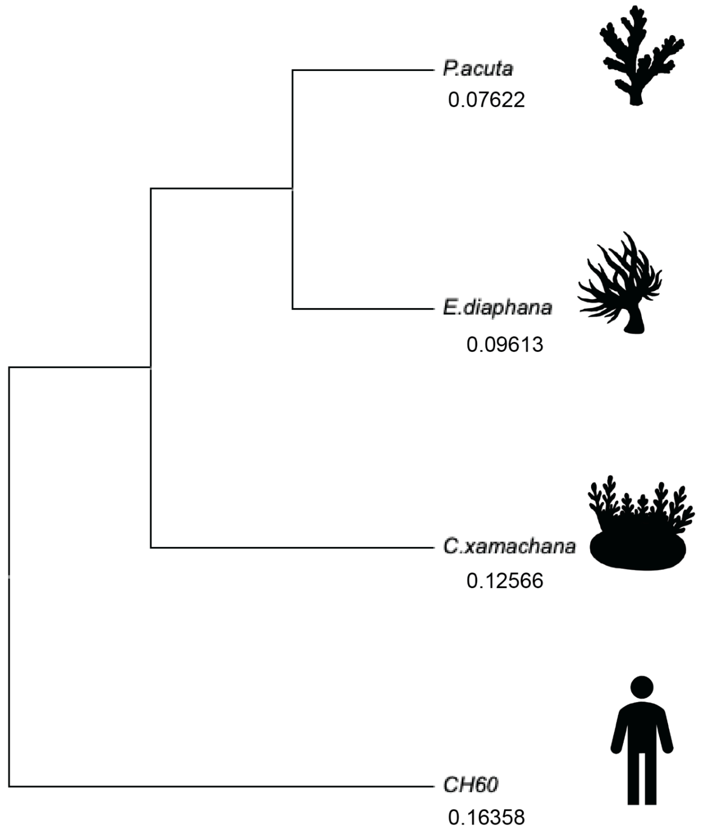


**Figure S5. Phylogenetic placement of cnidarian HSP60 proteins.** Rooted tree based on HSP60 (chaperonin-60) protein sequences from *P. acuta* (stony coral), *E. diaphana* (sea anemone), and *C. xamachana* (the upside-down jellyfish), with human HSP60/HSPD1 (CH60) used as the outgroup.

A

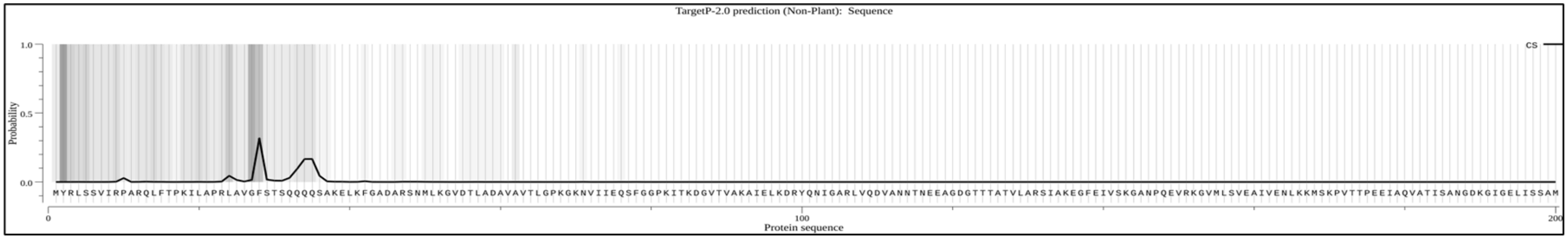


**Sequence**
**Prediction:** Mitochondrial transfer peptide
CS pos: 28-29. VGF-ST. Pr: 0.3169

| Protein type | Other | Signal peptide | Mitochondrial transfer peptide |
|---|---|---|---|
| **Likelihood** | 0.0089 | 0.0002 | 0.9909 |

B

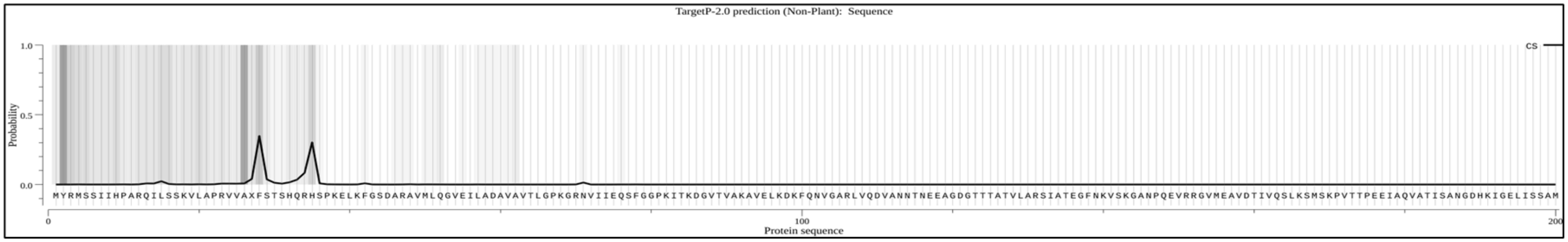


**Sequence**
**Prediction:** Mitochondrial transfer peptide
CS pos: 28-29. AXF-ST. Pr: 0.3479

| Protein type | Other | Signal peptide | Mitochondrial transfer peptide |
|---|---|---|---|
| **Likelihood** | 0.02 | 0 | 0.9799 |

C

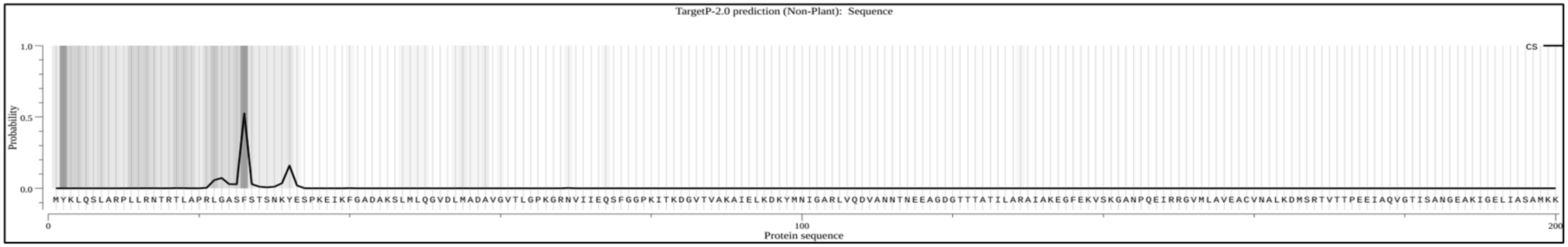


**Sequence**
**Prediction:** Mitochondrial transfer peptide
CS pos: 26-27. ASF-ST. Pr: 0.5249

| Protein type | Other | Signal peptide | Mitochondrial transfer peptide |
|---|---|---|---|
| **Likelihood** | 0.0042 | 0.0001 | 0.9957 |

**Figure S6. TargetP-2.0 outputs confirming mitochondrial localization.** Generated graphs confirming the location and confirmation of candidate sequences for mitochondrial HSP60 in (A) *P. acuta*, (B) *E. diaphana*, and (C) *C. xamachana,* as well as a probability table detailing the likelihood of mitochondrial signaling peptide with predicted cleavage sites.

**S7. Table showing alignment of candidate sequences.**

Table comparing the sequence alignment of candidate HSP60 protein sequences within *P. acuta* (stony coral), *E. diaphana* (sea anemone), and *C. xamachana* (upside-down jellyfish) against the known human HSP60 (chaperonin-60) sequence (P10809).

| Organism | Candidate sequence (source) | Length | Score | E-value | Identities | Positives | Gaps |
|---|---|---|---|---|---|---|---|
| *P. acuta* | TCONS_0030188 (Vidal-Dupiol et al, 2020) | 596 | 803 bits (2073) | 0 | 401/564 (71%) | 474/564 (84%) | 10/564 (1%) |
| *E. diaphana* | P18687 (Reef Genomics) | 586 | 808.52 (2087) | 0 | 394/566 (69.6%) | 480/566 (84.8%) | 10/566 (1.8%) |
| *C. xamachana* | Casxa1\|9735 (JGI) | 583 | 967 bits (2499) | 0 | 513/561 (91%) | 541/561 (96%) | 0/561 (0%) |

**S8. Table showing epitope alignments among candidate sequences.**

Table comparing the results generated from the epitope sequence of human HSP60 (p10809) epitope (amino acids 383-419) aligned with candidate HSP60 sequences in *P. acuta* (stony coral), *E. diaphana* (sea anemone), and *C. xamachana* (upside-down jellyfish).

| Organism | Candidate sequence (source) | Length | Score | E-value | Identities | Positives | Gaps |
|---|---|---|---|---|---|---|---|
| *P. acuta* | TCONS_0 030188 (Vidal-Dupi ol et al, 2020) | 37 | 68.2 bits (165) | $3x10^{-20}$ | 32/37 (86%) | 37/37 (100%) | 0/37 (0%) |
| *E. diaphana* | P18687 (Reef Genomics) | 37 | 68.6 bits (166) | $2x10^{-20}$ | 32/37 (86%) | 37/37 (100%) | 0/37 (0%) |
| *C. xamachana* | Casxa\|973 5 (JGI) | 37 | 68.6 bits (166) | $2x10^{-20}$ | 33/37 (89%) | 37/37 (100%) | 0/37 (0%) |
